# Shifts in temperature influence how *Batrachochytrium dendrobatidis* infects amphibian larvae

**DOI:** 10.1101/165985

**Authors:** Paul W. Bradley, Michael D. Brawner, Thomas R. Raffel, Jason R. Rohr, Deanna H. Olson, Andrew R. Blaustein

**Author notes:** **Author contribution statement** ARB, JRR, and TRR originally formulated the idea. PWB designed the experiment and developed the methodology. PWB performed the experiment. PWB and MDB performed the molecular analysis. PWB and TRR performed the statistical analyses. ARB, JRR, TRR, and DHO obtained funding. PWB wrote the manuscript and other authors provided editorial advice.

## Abstract

Many climate change models predict increases in mean temperature, and increases in frequency and magnitude of temperature fluctuations. These potential shifts may impact ectotherms in several ways, including how they are affected by disease. Shifts in temperature may especially affect amphibians, a group with populations that have been challenged by several pathogens. Because amphibian hosts invest more in immunity at warmer than cooler temperatures and parasites may acclimate to temperature shifts faster than hosts (creating lags in optimal host immunity), researchers have hypothesized that a temperature shift from cold-to-warm might result in increased amphibian sensitivity to pathogens, whereas a shift from warm-to-cold might result in decreased sensitivity. Support for components of this climate-variability based hypothesis have been provided by prior studies of the fungus *Batrachochytrium dendrobatidis* (Bd) that causes the disease chytridiomycosis in amphibians. We experimentally tested whether temperature shifts before Bd exposure alter susceptibility to Bd in the larval stage of two amphibian species – western toads (*Anaxyrus boreas*) and northern red legged frogs (*Rana aurora*). Both host species harbored elevated Bd infection intensities under constant cold (15° C) temperature in comparison to constant warm (20° C) temperature. Additionally, both species experienced an increase in Bd infection abundance when shifted to 20° C from 15° C, compared to a constant 20° C but they experienced a decrease in Bd when shifted to 15° C from 20° C, compared to a constant 15° C. These results are in contrast to prior studies of adult amphibians that found increased susceptibility to Bd infection after a temperature shift in either direction, highlighting the potential for species and stage differences in the temperature-dependence of chytridiomycosis.

## Introduction

Climate change represents one of the greatest challenges to biodiversity and conservation because it might compromise ecosystem functions worldwide. Changes in climate have affected plant-animal interactions, predator-prey interactions and disease dynamics (Lafferty 2009, Rohr *et al*. 2011, Sheldon, Yang & Tewksbury 2011, Garcia *et al*. 2014). Changes to annual or seasonal mean temperatures often are used to predict climate-change-induced effects on disease risk (Paaijmans, Read & Thomas 2009, Paaijmans *et al*. 2010). However, many climate change models also predict increases in the frequency and magnitude of extreme weather events and increases in temperature variability at monthly to weekly timescales (Easterling *et al*. 2000, Meehl & Tebaldi 2004, Schar *et al*. 2004, Paaijmans *et al*. 2010, Rummukainen 2012). Yet few studies have investigated how increases in temperature variability affect disease dynamics despite the likelihood that such variability might differentially affect hosts and pathogens (Paaijmans *et al*. 2010, Ben-Horin, Lenihan & Lafferty 2012, Raffel *et al*. 2013, Bannerman & Roitberg 2014, Luis *et al*. 2014, Raffel *et al*. 2015). Ectotherms, such as amphibians, are particularly sensitive to climate change (Blaustein *et al*. 2010, Lawler *et al*. 2010, Shoo *et al*. 2011, Li, Cohen & Rohr 2013) and are experiencing disease-associated population declines and extinctions worldwide (Stuart *et al*. 2004, McCallum 2007, Rohr *et al*. 2008, Wake 2012), making them an ideal group to investigate the relationship between temperature shifts and disease risk.

Chytridiomycosis is an emerging infectious disease of amphibians caused by the aquatic chytrid fungal pathogens *Batrachochytrium dendrobatidis* (Bd) and *B. salamandrivorans* (Longcore, Pessier & Nichols 1999, Martel *et al*. 2013). Bd is widespread globally (Liu, Rohr & Li 2013, Olson *et al*. 2013) and is associated with worldwide amphibian population declines (Stuart *et al*. 2004, Skerratt *et al*. 2007). Moreover, models based on IPCC climate futures predict that Bd will shift to higher latitudes and altitudes due to increased environmental suitability in those regions under climate change, thus potentially affecting additional amphibian populations (Xie, Olson & Blaustein 2016).

The negative effects of Bd infection are more pronounced in post-metamorphic stages, often leading to death (Blaustein *et al*. 2005, Garner *et al*. 2009, Gervasi *et al*. 2013, Gervasi *et al*. 2017). In larvae, Bd infection can cause host mortality in some species (Blaustein *et al*. 2005, Garner *et al*. 2009). However the infection is localized to keratinized larval mouthparts, (Marantelli *et al*. 2004, McMahon & Rohr 2015) often resulting in sublethal effects including inhibited foraging capacity, reduced growth and development, altered predator avoidance, or changes to other behaviors (Han, Bradley & Blaustein 2008, Venesky, Parris & Storfer 2010, Buck *et al*. 2012, Gervasi *et al*. 2013). Additionally, larvae of many species are important members of aquatic communities and alterations to larval feeding have the potential to cascade through the aquatic ecosystem (Alford 1989, Brönmark, Rundle & Erlandsson 1991, Lamberti *et al*. 1992, Kupferberg 1997).

Temperature is considered one of the most important environmental factors driving chytridiomycosis (Drew, Allen & Allen 2006, Bosch *et al*. 2007, Daskin, Alford & Puschendorf 2011, Forrest & Schlaepfer 2011, Voyles *et al*. 2017). Bd is non-linearly sensitive to temperature with an optimal growth range in culture between 17° and 25° (Piotrowski, Annis & Longcore 2004, Rohr & Raffel 2010, Raffel *et al*. 2013) and a temperature-dependent generation time of 4 to 10 days (Woodhams *et al*. 2008). The upper thermal limit for Bd growth in culture is between 25°C and 28°C, with Bd mortality occurring above 30°C (Longcore, Pessier & Nichols 1999, Piotrowski, Annis & Longcore 2004). Bd has been shown to be reliably cleared from multiple amphibian species by extended exposure to 30°C (McMahon *et al*. 2014). Its lower thermal limit is below 4°C (Piotrowski, Annis & Longcore 2004). Additionally, life history strategies of the pathogen can be altered by environmental temperature, where colder temperatures can cause Bd zoosporangia to develop and mature more slowly (Voyles *et al*. 2012), but produce more and longer-lived zoospores overall (Hyatt *et al*. 2007, Woodhams *et al*. 2008).

Because physiologies of both the host and pathogen are strongly influenced by environmental temperature, climate change has been used to explain several major Bd outbreaks and amphibian population declines, (reviewed in Li, Cohen & Rohr 2013, Rohr *et al*. 2013). Yet, the host and pathogen are not expected to share a uniform response to a given temperature (Brown *et al*. 2004, Paull, LaFonte & Johnson 2012, Rohr *et al*. 2013), and thermal responses measured in constant-temperature artificial environments might not reflect organism responses in more realistic variable-temperature environments. Providing evidence of the lack of a uniform response between Bd and amphibians to temperature shifts, Rohr and Raffel (2010) found a strong correlation between elevated month-to-month temperature variability and Bd-associated amphibian population declines of *Atelopus* spp. across Central and South America. Further support of the relationship between chytridiomycosis and temperature variation has been provided by laboratory studies. In one study, Cuban treefrogs (*Osteopilus septentrionalis*) displayed reduced resistance to Bd infection when exposed to random daily temperature fluctuations or when exposed to a temperature decrease after acclimation to a warmer temperature (Raffel *et al*. 2013). Similar results were obtained in newts (*Notophthalmus viridescens*) exposed to Bd, except both decreases and increases in temperature were associated with elevated Bd abundance relative to abundances at constant temperatures (Raffel *et al*. 2015).

The potential for temperature variability to increase disease severity in amphibians was first postulated by Raffel *et al*. (2006) and has subsequently been referred to as the “climate variability hypothesis” (Rohr & Raffel 2010). This hypothesis posits that parasites acclimate to the new temperature more rapidly than their hosts, leading to lags in host acclimation following a temperature shift that could make hosts more susceptible to infection (Raffel *et al*. 2013). This hypothesis assumes that: 1) pathogens acclimate to the new temperature faster than the host because of their relatively smaller size and higher metabolic rate (Gillooly *et al*. 2001, Raffel *et al*. 2013); and 2) both host and parasite acclimation responses lead to increased performance at the new temperature, in accordance with the “beneficial acclimation hypothesis” of thermal biology (Angilletta 2009). However, Raffel *et al*. (2006) also pointed out potential complexities in acclimation of the ectotherm immune system that might lead to alternative predictions.

According to the “*lag effect*” hypothesis (Raffel *et al*. 2006), changes in levels of temperature-dependent immune parameters might simply lag behind environmental temperature shifts (Fig. 1) because it takes time to produce necessary, or remove unnecessary, immune cells from the host. For example, amphibians are expected to require more immune cells at warmer temperatures to fight off faster-growing pathogens (Maniero & Carey 1997), and lags in production of new immune cells could lead to sub-optimal immunity following a temperature increase (Raffel *et al*. 2006). Conversely, the amphibian immune system is expected to be down-regulated following a temperature decrease (Macela & Romanovsky 1970), with the removal of mature white blood cells determined by the rate of their respective half-lives (DeSantis & Strauss 1997, Janeway 2008). A lag in this process might lead to a brief period of elevated immune responsiveness relative to an already cold-acclimated host. Thus, the “lag effect” hypothesis predicts the opposite effect from the “climate variability hypothesis” following a temperature decrease, at least on a short timescale. These mechanistic hypotheses are not mutually exclusive, and it is unclear which effects might be more important for a given host-parasite combination.

**Fig 1.**
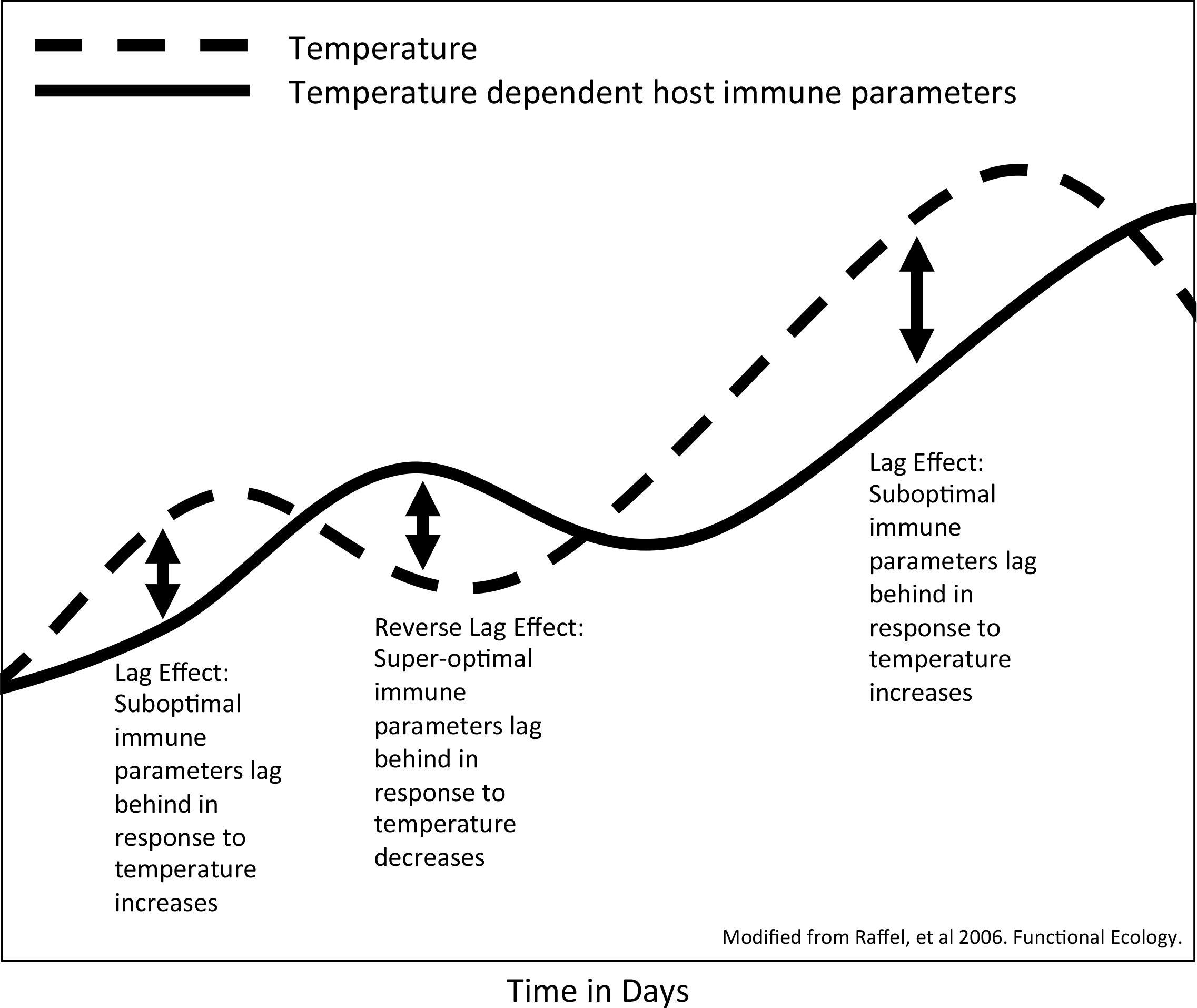
Hypothesized lag effect showing the relationship between fluctuating temperatures (over days to weeks) and the optimal levels of a hypothetical temperature-dependent host immune parameter. The immune parameter follows and lags behind temperature changes – resulting in periods of a compromised immune status after a temperature increase, and resulting in an over-active (or unnecessarily costly) immune status after a temperature decrease. Modified from Raffel *et al*. (2006).

We tested the general prediction that an amphibian shifted to a new temperature before Bd exposure would respond to infection differently than a host already acclimated to the exposure temperature. We postulated that the direction of the effect would depend upon the direction of the temperature shift, in accordance with the “lag effect” hypothesis of Raffel *et al*. (2006). Given the differences in size between the host and the pathogen, and associated physiological process rate differences, we assumed Bd would physiologically respond to the temperature shift faster than the host, such that an idealized host-immune response to Bd exposure would temporarily lag behind the temperature shift. Thus, we predicted that a temperature shift from cold-to-warm would result in an *increase* in susceptibility to Bd exposure, whereas a temperature shift from warm-to-cold would result in a *decrease* in susceptibility to Bd exposure. To test these predictions, we quantified susceptibility to Bd by measuring infection abundance after exposure to the pathogen.

## Materials and Methods

To examine the how temperature shifts may alter larval amphibian infection dynamics, we selected two species of amphibian hosts, the northern red legged frog (*Rana aurora*) and the western toad (*Anaxyrus boreas*). Both species have been observed in the field with Bd infections (Pearl *et al*. 2007, Muths, Pilliod & Livo 2008, Piovia-Scott *et al*. 2011) and both species are susceptible to chytridiomycosis (Han, Bradley & Blaustein 2008, Gervasi *et al*. 2013). To ensure that the animals used in our experiment were not previously infected with Bd, amphibians were collected as eggs from natural oviposition sites. Red legged frog eggs were collected from a permanent pond located near Florence, Oregon, USA (Lincoln County, elevation 12 m; latitude/longitude: 44.088/-124.123) in the Oregon Coast Range on 11-Feb-2012. Western toad eggs were collected from Little Three Creeks Lake (Deschutes County, elevation 2,000 m; latitude/longitude: 44.009/-121.643) in the Cascade Range on 9-Jul-2011. Immediately after collection, eggs were transported to a laboratory at Oregon State University where they were maintained at 14° C, under a 12-12 photoperiod in 40-liter aquaria filled with dechlorinated water. Upon hatching, larvae were maintained at a density of approximately 200 individuals per aquarium and fed *ad libitum* a mixture of Tetramin fish food and ground alfalfa pellets (1:3 ratio by volume). Water was changed every seven days. The 40-day trials for each species were not run concurrently, but identical protocols were used for both species and both trials consisted of individuals of identical larval stage (Gosner stage 26).

### Acclimation Period

Independent trials for each host species began with a 20-day acclimation period with 80 (Gosner stage 26) larvae randomly selected, and individually placed into 80 plastic 500-mL containers where they were housed for the duration of the acclimation period and experiment. Each container was filled with 14° C dechlorinated water and covered with a lid to help maintain water temperature and limit evaporation. Each container had 2-mm diameter holes drilled between the water line and the lid to allow air circulation into the container. Pairs of containers were then placed within 40 individual temperature-controlled chambers (to ensure independent replication of the temperature treatments) that were set at 15° C to avoid cold-shocking the larvae. Each temperature-controlled chamber was independently controlled via its own thermostat and the interior measured approximately 37 cm deep x 21 cm wide x 13 cm in height. Half of the 40 temperature-controlled chambers were then randomly selected to begin the acclimation period at 20° C (warm treatment) and the other half were kept at 15° C (cold treatment). The placement of temperature chambers within the laboratory was randomized, as was the placement of 500-mL containers within each temperature chamber.

### Temperature Shifts

On day 20 of the experiment, half of the temperature chambers in each of the two acclimation temperatures (15° C and 20° C) were randomly selected to undergo a temperature shift, either from 20° to 15° C or from 15° C to 20° C. The other half of the temperature chambers underwent no shift in temperature. Thus, each of the temperature chambers was subjected to one of four temperature treatments: a constant 15° C (cold) throughout the experiment; a constant 20° C (warm) throughout the experiment; a temperature shift from 15° C to 20° C (cold-to-warm); or a temperature shift from 20° C to 15°C (warm-to-cold).

### Bd exposure

On day 24, one of the two 500-mL containers within each temperature-controlled chamber was randomly selected to undergo a Bd-exposure treatment and the other was selected as a control. Larvae in the Bd-exposure treatment were exposed to a single inoculate of Bd strain JEL 274, which was grown in pure culture on 1% tryptone agar in 10-cm diameter Petri dishes. This Bd strain was selected as it is one of the more virulent strains associated with major amphibian populations declines (Rosenblum *et al*. 2013). The Petri dishes were inoculated with liquid culture 10 days before the start of the experiment and incubated at 15° C. To harvest the zoospores, 10 plates were flushed with 15 mL of 15° C dechlorinated water and remained undisturbed for 10 minutes. The plates were scraped with a rubber spatula to release the zoospores and sporangia adhering to the agar. The inoculum from each plate was then pooled in a beaker and the number of moving zoospores was determined using a hemocytometer. After quantifying the zoospore concentration, the inoculum was diluted to 10,000 zoospores/mL. Individuals in the Bd-exposed treatments were exposed to 10 mL of inoculum transferred into the 500-mL container housing an individual larva. Control individuals were exposed to 10 mL of sham inoculum lacking the Bd culture (made from 1% tryptone sterile agar plates following the same methods), similarly transferred into the 500-mL container housing each larva. Thus, the individual larva underwent their exposure treatment on day 24, four days after the water temperature shift for chambers in the two temperature shift treatments.

During the 40-d trial, survival and metamorphic status were checked daily. Water for each 500-mL container within the temperature chambers was changed every 12 days and consisted of dechlorinated water of the same temperature (15° C and 20° C). Individuals that survived until the end of the trial (i.e., day 40) were euthanized in a 2% solution of MS-222, and then preserved in 95% ethanol. Individuals that reached metamorphosis (Gosner stage 42: emergence of forelimbs) were euthanized, measured, and preserved as previously described.

### Determining infection status

We used quantitative polymerase chain reaction (qPCR) to determine infection status and quantify Bd-infection intensity of all individuals in the Bd-exposure treatments. Additionally, we investigated Bd-infection status in eight randomly selected control individuals per species. To sample the individuals for Bd, we extracted whole mouthparts of the larvae using sterile dissection scissors. We conducted qPCR using an ABI PRISM 7500 sequencer (Applied Biosystems) according to the methods of Boyle *et al*. (2004) except that we used 60 µL of Prepman Ultra (Applied Biosystems, Carlsbad, California, USA), instead of the 40 µL in the DNA extraction. All samples were run in triplicate and averaged.

### Statistical Analyses

Each temperature-controlled chamber was an experimental unit (whole plot) and the pairs of containers within each chamber acted as subplots. The whole plots were subjected to one of four temperature regimes consisting of a Bd-exposure temperature combined with a temperature shift status (constant cold, constant warm, shifted to cold, and shifted to warm). Further, subplots were subjected to one of two exposure treatments (Bd exposed and Bd unexposed).

Survival was compared between temperature treatments for western toad larvae with a Cox proportional hazards model (Cox 1972) using TIBCO Spotfire S+ version 8.1. The model consisted of the main effects of the temperature treatment, temperature shift status (constant versus shifted), and an interaction between the two variables. Due to losses of western toad larvae prior to the application of the exposure treatment, we lacked the power to statistically compare survival in western toad larvae between the Bd exposure treatments

Bd infection abundance (Bd genomic equivalents) among temperature treatments and between host species was analyzed using R version 3.11. We used a zero-inflated negative-binomial generalized linear model (function ‘zeroinf’ in package ‘pscl) as described by Raffel *et al*. (2010), which includes a zero-inflation component that models infection status as a binomial process (binomial distribution with a logit link) and a count component that models infection intensity as a negative binomial process (negative binomial distribution with a log link). Our full model investigated the effects of all of the explanatory variables including host species, exposure temperature, temperature shift status, and all two- and three-way interactions on Bd (*Batrachochytrium dendrobatidis*) abundance. Interpretation of this analysis required further reduced models to investigate the effect of exposure temperature and temperature shift for each species (species model) and the effect of temperature shift for each Bd-exposure temperature and host species combination (Bd-exposure temperature model).

## Results

Survival differences were not detected between exposure temperatures (Cox, Z = -1.099, *p* = 0.27) or temperature shift status (Cox, Z = -0.277, *p* = 0.78) in Bd-exposed western toad larvae. We were unable to detect survival differences in red legged frog larvae, as only one individual larva experienced mortality after application of the exposure treatment (Table S1).

### Infection Abundance

We detected a host species by temperature shift interaction (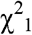 = 3.83, *p* = 0.050; Table S2) and a Bd-exposure temperature by temperature shift interaction (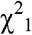 = 7.50, *p* = 0.006; Table S2). We investigated these interactions with reduced models to investigate effects on Bd abundance at the levels of species and exposure temperature.

Red legged frog larvae had higher Bd abundance when they were exposed to infection at 15° C when compared to 20° C (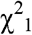 = 3.88, *p* = 0.049; Fig. 2). The main effect of temperature shift was marginally significant in the reduced species model analysis (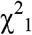 = 3.50, *p* = 0.061), but there was a significant effect of temperature shift for individuals exposed at 20° C in the reduced Bd-exposure model (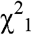 = 5.7, *p* = 0.017), with individuals shifted from 15° C to 20° C having higher Bd abundance than red legged frog larvae experiencing constant 20° C (Fig. 2). In contrast, there was no evidence that a temperature shift influenced Bd infection when red legged frog larvae were exposed to Bd at 15° C (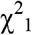 = 0.6, *p* = 0.4; Fig. 2). There was no statistically significant interaction between exposure temperature and temperature shift for red legged frog larvae (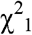 = 2.4, *p* = 0.13).

**Fig 2.**
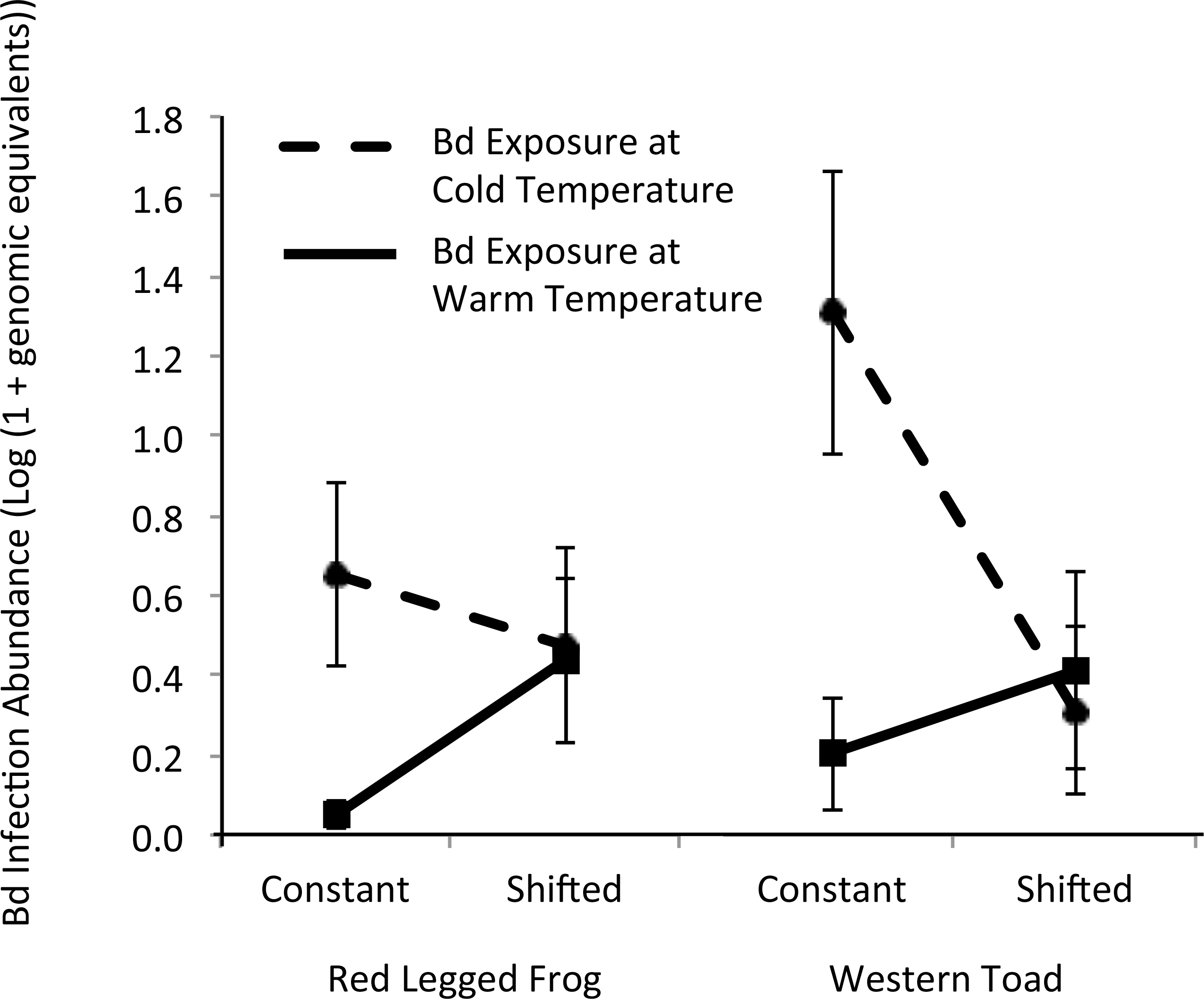
Mean *Batrachochytrium dendrobatidis* (Bd) infection abundance (± SE) measured at death, or at euthanasia 16-days after Bd exposure, in both western toad (*Anaxyrus boreas*) larvae and red legged frog (*Rana aurora*) larvae from Oregon, USA, and between the two temperatures at the time of Bd-exposure (cold [15° C] versus warm [20° C]) and between larvae having experienced either a constant or shifted temperature. Bd infection abundance is quantified as the log (1 + Bd genomic equivalents) per excised larval mouthparts of all individuals exposed to the pathogen.

We detected an interactive effect of exposure temperature and temperature shift on Bd abundance in western toad larvae (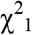 = 5.2, *p* = 0.023). This was driven by elevated Bd abundance in individuals under the constant 15° C temperature when compared to individuals that experienced a temperature shift from 20° to 15° C, but no evidence of an effect of shifting temperature from 15° C to 20° C (Fig. 2). There were no main effects of exposure temperature (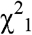 = 0.50, *p* = 0.5) or temperature shift (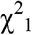< 0.01, *p* = 0.9) on Bd abundance in western toad larvae. Further, when investigating the exposure temperatures individually in the reduced Bd-exposure model, there was no evidence that a temperature shift influenced Bd infection in western toad larvae after exposure to Bd at 15° C (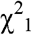 = 3.4, *p* = 0.066) or 20° C (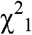 = 2.5, *p* = 0.11).

We failed to find evidence that the two host species differed in response to exposure to the pathogen, leading us to conclude that general patterns for both species were similar (Fig. 2). Both species experienced an increase in Bd abundance when shifted to 20° C compared to a constant 20° C, and both generally experienced a decrease in Bd abundance when shifted to 15° C compared to a constant 15° C. Additionally, both host species experienced elevated Bd abundance in the constant 15º C treatment when compared to the constant 20° treatment.

All red legged frog individuals survived until the end of the experiment but a number of western toad individuals died or metamorphosed earlier (Table S2). We therefore assessed the possibility that the timing of Bd sampling or the proximity of a larva to metamorphosis might drive observed patterns of Bd abundance in western toads. The model for Bd abundance on western toads was not significantly improved by adding either a variable coding whether individuals were near metamorphosis when sampled (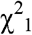 = 4.00, *p* = 0.150) or a covariate indicating the sampling date (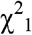 = 3.33, *p* = 0.068). Furthermore, neither variable qualitatively changed the contribution of exposure temperature or temperature shift status to the model. Therefore, we omitted both covariates from the final model for western toads.

## Discussion

Our results suggest that Bd infection dynamics in larval amphibians can be affected by a shift in water temperature before host exposure to the pathogen, and that the direction of temperature shift determines the outcome of Bd exposure. Similar patterns were observed for the two host species when comparing individuals exposed to constant versus shifted temperatures. A shift from the warm temperature to the colder temperature was associated with a significant decrease in Bd abundance in western toad larvae and no significant decrease in red legged frog larvae. Likewise, a shift from the cold temperature to the warmer temperature significantly increased Bd abundance in red legged frog larvae and had no significant effect in western toad larvae. Importantly, we detected the effects of temperature shifts despite the host having a four-day head start on acclimating to the Bd exposure temperature relative to the pathogen. This suggests that we are likely underestimating the strength of these effects and that their magnitudes might have been larger if the host and pathogen experienced the shifts concurrently.

Amphibian species do not all respond similarly to a given Bd exposure. Species-level differences in host tolerance to Bd infections have been well documented under controlled laboratory conditions (Searle *et al*. 2011, Gervasi *et al*. 2017). Under natural conditions, pathogen tolerance within a species may be affected by biotic factors such as inter- and intra-specific interactions, proximity to metamorphosis, or life stage (Parris & Cornelius 2004, Rachowicz & Vredenburg 2004, Blaustein *et al*. 2005, McMahon & Rohr 2015) or abiotic factors such as temperature, season, or resource availability (Berger *et al*. 2004, Raffel *et al*. 2010). For some susceptible host species, temperature-shift induced changes in Bd abundance might alter the outcome of infection by either pushing *Bd* abundance over or under a tolerance threshold. Such changes in relation to pathogen abundance and pathogen tolerance may result in altering the strength of negative effects of Bd infection. For example, temperature shifts in synergy with Bd infection may result in either positive or negative effects on growth and development rates, foraging efficiency, or predator avoidance (Parris & Cornelius 2004, Parris, Reese & Storfer 2006, Venesky, Parris & Storfer 2010, Venesky, Wassersug & Parris 2010).

We hypothesized that hosts exposed to a shifted temperature would respond to infection differently than hosts exposed to a constant temperature, and under the framework of the “lag effect” hypothesis (Raffel *et al*. 2006, Rohr & Raffel 2010), the direction of the temperature shift would differentially affect infection severity. We predicted that a temperature shift from cold-to-warm would leave hosts in a temporarily immune-compromised state and result in an elevated Bd abundance after exposure when compared to hosts exposed to a constant warm temperature. Conversely, we predicted that a temperature shift from warm-to-cold would provide hosts with a temporarily elevated-immune responsiveness and result in a decrease in Bd abundance after exposure when compared to hosts exposed to a constant cold temperature. Our results were consistent with predictions of the “lag effect” hypothesis, and were generally consistent with previous studies showing that a shift in temperature influences Bd infection in postmetamorphic amphibians (Raffel *et al*. 2013, Raffel *et al*. 2015). In particular, our finding of decreased resistance to infection following a temperature increase (relative to warm-acclimated individuals) mirrored a laboratory study of post-metamorphic red-spotted newts (*Notophthalmus viridescens*), where juvenile newts exhibited decreased Bd resistance following a shift from 15° C to 25° C (Raffel *et al*. 2015). These findings of fluctuating temperature effects on Bd infection across four anuran taxonomic groups and life-stages suggest that effects of temperature shifts and Bd-related chytridiomycosis susceptibility might be widespread within amphibians. However, our finding of increased resistance to Bd infection following a temperature decrease (relative to cold-acclimated individuals) was opposite the pattern observed in red-spotted newts and Cuban treefrogs (Raffel *et al*. 2013, Raffel *et al*. 2015) These contrasting results suggests that there are important among-taxa or among-stage differences in the underlying mechanisms driving the effects of temperature fluctuation on Bd infection; whereas our results in premetamorphic life-stage of western toads and red legged frogs are consistent with the “lag effect” hypothesis, results of similar studies investigating post-metamorphic red-spotted newts and Cuban treefrogs support the “climate variability hypothesis.”

We observed differences in Bd abundance on our two amphibian species at the two constant temperature treatments. Higher Bd abundances were observed for both host species under the constant cold temperature treatment compared to the constant warm temperature treatment. These results are consistent with previous experiments that showed increased Bd abundance (Raffel *et al*. 2015) and Bd-induced mortality (Kilpatrick, Briggs & Daszak 2010, Murphy, St-Hilaire & Corn 2011, Raffel *et al*. 2015) associated with lower temperatures. This is despite Bd growing best in culture at about 23° C, which is much closer to the warm than cold temperature treatments in this experiment (Piotrowski, Annis & Longcore 2004, Woodhams *et al*. 2008). This might be because the larval immune response to Bd infection increases with increasing temperatures at a faster rate than the infectivity or growth rate of Bd (Raffel *et al*. 2013), or alternatively because of the differences between the growth rate of Bd in culture compared to the growth rate on host tissue (Venesky *et al*. 2013). Our results provide further evidence to suggest patterns of Bd growth in culture differ from patterns of Bd growth on a host and that it is important to assess the host-parasite interaction when predicting effects of climate and climate change on disease risk.

Alternatively, differences in Bd abundance between the two constant temperature treatments might be due to temperature effects on the pathogen rather than the host (Woodhams *et al*. 2008, Voyles *et al*. 2012). The Bd was cultured at 15° C; it is possible that the temperature shift experienced by the pathogen in the warm exposure treatment caused the depressed Bd abundances observed in both host species compared to the elevated Bd abundance in the cold exposure temperature treatment. A decrease in temperature may cause an increase in the number of Bd zoospores released from zoosporangia (Hyatt *et al*. 2007, Woodhams *et al*. 2008), however the effect of a similar increase in temperature on Bd physiology is unclear.

In conclusion, our results provide additional evidence for climate variability affecting Bd infection in amphibians but suggest important among-taxa differences in the directionality of these effects. Our finding of increased host resistance to infection following a temperature decrease is consistent with the “lag effect” hypothesis of Raffel *et al*. (2006) but contradicts components of the “climate variability hypothesis”, which has been proposed as an explanation for patterns of Bd-associated amphibian population declines (Rohr & Raffel 2010, Raffel *et al*. 2013, Raffel *et al*. 2015). Our study highlights the complexity that temperature plays in determining the outcome of Bd-amphibian interactions and the role that a fluctuating temperature might play in altering these interactions. Furthermore, this study increases the diversity of amphibian species and stages that have been shown to exhibit thermal acclimation effects on disease, and the broad generality of this pattern across four disparate taxa suggests that fluctuating-temperature effects on amphibian infection may be widespread. Accurately predicting the effects of global climate change on infectious diseases, such as chytridiomycosis will require further understanding of how infectious agents respond to heterogeneity in temperatures and temperature fluctuations.

## Acknowledgments

All applicable institutional and national guidelines for the care and use of animals were followed; this research was conducted under Oregon State University IACUC animal care and use permit 3917. Collection of amphibian eggs was approved by the Oregon Department of Fish and Wildlife (Oregon Scientific Taking Permit #006-12 issued to ARB). We thank S. Bauer, E. Davis, E. Hunt, A. Koosman, B. Meyers, M. Ouspenskaya, E. Peseke, V. Raffeale, and C. Rains for their help performing the experiment, K. Boersma for her help with the experimental design, and E. Boersley for her support and assistance. Additionally we thank J. Spatafora, V. Weis, and the Center for Genome Research and Biocomputing at Oregon State University for providing laboratory space for qPCR. This research was supported by grants from the National Science Foundation (EF-1241889), National Institutes of Health (R01GM109499, R01TW010286), U.S. Department of Agriculture (NRI 2006-01370, 2009-35102-0543), and U.S. Environmental Protection Agency (CAREER 83518801) to JRR and NSF grant IOS 1121529 to TRR. Support was provided by the U.S. Forest Service Pacific Northwest Research Station, Corvallis, Oregon to DHO.

## Supporting Information

Additional supporting information may be found in electronic supplementary material for this article.

